# Directed evolution of compact synthetic promoters via AlphaGenome and genetic algorithms

**DOI:** 10.64898/2026.06.28.735069

**Authors:** Linxiao Nie

## Abstract

Compact tissue-specific promoters are highly desirable for gene therapy because viral vectors possess limited packaging capacity. However, existing promoter engineering strategies rely primarily on rational design or de novo sequence generation and lack efficient approaches for compressing long native promoters while preserving regulatory specificity. Although genome foundation models have substantially improved sequence-to-function prediction, they have not been effectively translated into computational platforms for promoter engineering.

Here, we present VirEvo, a computational promoter engineering framework that integrates a virtual dual-luciferase assay (VirDLA), genome-foundation-model-guided genetic evolution, and an orthogonal Pan-Tissue Consistency Filter (PTCF). VirDLA introduces an internal-reference normalization strategy inspired by dual-luciferase reporter assays, enabling relative comparison of promoter activity across tissues without retraining AlphaGenome. Guided by these normalized activity scores, VirEvo iteratively optimizes promoter selectivity, off-target activity, and sequence length.

Using the human p16^INK4a^ promoter as a proof of concept, VirEvo evolved a compact synthetic promoter, SRP2M, of only 398 bp, representing an 85.9% reduction in sequence length. Experimental validation using dual-luciferase reporter assays in senescent IMR90 fibroblasts demonstrated that SRP2M retained 77% of wild-type senescence selectivity while reducing basal leakage to 52% of the wild-type level.

Together, these results demonstrate the feasibility of genome-foundation-model-guided promoter engineering. VirEvo provides a generalizable framework for designing compact tissue-specific regulatory elements and extends the application of genome foundation models from functional prediction to synthetic regulatory engineering.

## INTRODUCTION

Gene therapy is highly dependent on safe, efficient, and specific gene delivery systems. Among these vectors, Adeno-Associated Virus (AAV) has emerged as one of the most widely used platforms in clinical applications due to its low immunogenicity and favorable tissue tropism. However, AAV possesses a limited effective packaging capacity; once the payload exceeds approximately 4.7 kb, its packaging efficiency and titer decline sharply^1^, approaching zero rapidly when the insert size slightly surpasses this threshold. Lentiviral vectors can accommodate approximately 10 kb of exogenous sequences ^2^ and achieve stable expression through genomic integration; however, they are associated with risks of insertional mutagenesis and relatively high production costs. Consequently, the limited packaging capacity of these vectors severely constrains the co-delivery of therapeutic genes and their essential regulatory elements.

Native tissue-specific promoters often occupy a significant portion of vector capacity due to their length. For instance, the liver-specific thyroglobulin promoter (>1.5 kb) and certain cardiac-specific promoters (e.g., the *α*-myosin heavy chain promoter >5 kb) are notably long. Concurrently, these promoters typically exhibit high basal expression ^2,3^, Consequently, increasing efforts have been devoted to developing compact synthetic promoters to improve gene delivery efficiency while maintaining tissue specificity and transcriptional activity ^4^.

Traditional rational design approaches attempt to address these issues by truncating or engineering native promoters; however, they face multiple challenges. These methods rely heavily on prior biological knowledge and experimentally characterized regulatory elements, making promoter development labor-intensive and difficult to systematically optimize. Many synthetic promoters generated through rational design have indeed been shortened ^3,5^, yet they still exhibit basal expression (leakiness) and struggle to maintain tissue or cell-type specificity. translating promoter performance from in vitro screening to in vivo applications remains a major challenge, as regulatory activity measured in cultured cells is often not predictive of in vivo performance ^6^. Furthermore,strong constitutive promoters, while offering high expression levels, lack the necessary tissue specificity; conversely, attempts to enhance specificity frequently compromise expression strength or inadvertently increase leakage.In recent years, deep learning models such as Enformer and Borzoi have made significant advances in predicting gene regulatory patterns ^7,8^. Enformer is trained on large-scale epigenetic data to predict gene expression and chromatin states from DNA sequences; Borzoi focuses on predicting cell-type- and tissue-specific RNA-seq coverage. However, these models are fundamentally predictive tools rather than iterative evolutionary frameworks designed for promoter length compression and functional optimization. If applied directly as fitness functions within genetic algorithms, they face severe resolution constraints: Enformer achieves a 128 bp resolution, while Borzoi is limited to 32 bp, making it difficult to accurately capture functional changes resulting from subtle sequence perturbations at the scale of hundreds of base pairs in synthetic promoters. In contrast, AlphaGenome, recently released by DeepMind, has achieved single-base-resolution predictions and represents a leading performance benchmark in this field ^9^. While suitable as a high-precision baseline model for assessing sequence fitness, it remains a predictive tool and lacks the evolutionary search mechanisms required for directed compression. Furthermore, although existing research has utilized GANs or diffusion models to generate synthetic promoters/enhancers, these approaches mostly focus on de novo design of strong constitutive promoters, lacking tissue specificity and rarely originating from long natural sequences for targeted compression optimization^10,11^. To our knowledge, no existing framework combines genome-foundation-model prediction, directed evolutionary search, and experimental validation for the compression of long native tissue-specific promoters.

p16^INK4a^ (CDKN2A) is a core marker of cellular senescence, with its promoter exhibiting specific activation in senescent cells; consequently, it has been widely employed to drive senolytic strategies. Previous studies^12^ demonstrated that clearing p16^+^ senescent cells can delay the progression of various age-related diseases and improve healthspan. However, the wild-type p16 promoter is exceptionally long at 2.6–2.8 kb ^13^, exhibits limited cell-type selectivity, and to date, there have been no reports of engineered compact designs. These characteristics make it an ideal proof-of-concept system for validating the balance between promoter compression and the retention of tissue specificity.

To address the lack of a framework capable of performing directed compression of long tissue-specific promoters while preserving regulatory specificity, we developed VirEvo, a computational promoter engineering platform that integrates a virtual dual-luciferase assay (VirDLA), genome-foundation-model-guided evolutionary search, and orthogonal candidate screening. By combining promoter selectivity, off-target activity, and sequence length into a unified optimization objective, VirEvo enables the iterative evolution of compact synthetic promoters directly from long native regulatory sequences. Genetic algorithms were selected as the search engine because promoter compression requires simultaneous exploration of both point mutations and large structural alterations—including insertion, deletion, duplication, and truncation—which naturally map onto genetic operators while allowing sequence length to vary throughout optimization.

As a proof-of-concept, we applied VirEvo to the human p16^INK4a^ promoter. Starting from the 2829 bp wild-type sequence, the framework evolved SRP2M, a synthetic promoter only 398 bp in length. Functional characterization using dual-luciferase reporter assays demonstrated that SRP2M retained substantial senescence-selective activity despite an 85.9% reduction in sequence length.

More broadly, VirEvo establishes a generalizable workflow for promoter compression and optimization that relies primarily on in silico evolution rather than large-scale experimental screening. Because the framework is largely independent of promoter identity and target tissue, it can in principle be adapted to other regulatory systems by replacing the starting promoter sequence and optimization context. These results suggest a potential route toward scalable computational design of compact tissue-specific regulatory elements for gene therapy and synthetic biology applications.

## METHODS

### 2.1 Design and Construction of the Virtual Dual-Luciferase Assay

To enable standardized comparison of regulatory activity across tissues and cellular contexts, we developed a Virtual Dual-Luciferase Assay (VirDLA). The framework is conceptually inspired by conventional dual-luciferase reporter assays, in which the signal generated by a test reporter is normalized against an internal reference reporter to reduce experimental variability. In VirDLA, the same principle is applied to genome-foundation-model predictions. By expressing activity as the ratio between a target reporter and an internal reference reporter, raw AlphaGenome outputs are transformed into a relative activity metric that is less sensitive to differences in sequence context, prediction baseline, and tissue-specific signal magnitude, thereby facilitating cross-context comparison.

#### 2.1.1 Reporter cassette design

To implement this strategy, a standardized virtual reporter cassette (Supplementary Figure 1)was constructed. The cassette consists of five functional modules arranged from upstream to downstream: a Variable Regulatory Region (VRR), a Target Reporter Module (TRM), an insulator element, an Internal Reference Promoter (IRP), and a Reference Reporter Module (RRM). The VRR serves as the regulatory sequence under investigation and may contain native promoters, synthetic promoters, engineered variants, or other regulatory elements. The TRM and RRM each contain a reporter coding sequence, a 3’ untranslated region, and a polyadenylation signal. The IRP constitutively drives expression of the RRM and provides the internal normalization signal required for activity calculation. To better match the sequence distribution encountered during AlphaGenome training, all non-coding regions of the cassette were humanized.

#### 2.1.2 Activity calculation

For each VirDLA evaluation, the candidate regulatory sequence was inserted into the VRR position of the reporter cassette to generate a complete virtual construct. The construct was then embedded at the center of a 1,048,576-bp sequence window and padded with neutral sequence prior to AlphaGenome inference.

AlphaGenome was subsequently used to predict strand-specific RNA-seq coverage across the entire cassette. Mean predicted RNA-seq coverage was calculated across the coding-sequence regions of both the Target Reporter Module (TRM) and the Reference Reporter Module (RRM), yielding the target reporter signal and reference reporter signal, respectively.

The normalized activity score for a given tissue or cellular context was defined as:

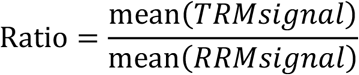

where TRM and RRM denote the predicted RNA-seq signals of the target and reference reporter modules, respectively. This ratio serves as the primary VirDLA activity metric and provides a relative measure of regulatory activity that is less sensitive to context-dependent variation in raw model outputs.

#### 2.1.3 External benchmarking using SHARPR-MPRA

For independent validation of the VirDLA framework, we used the SHARPR-MPRA K562 scale-up dataset ^14^, comprising 1,000 human genomic enhancer fragments (295 bp) with experimentally measured activity scores in K562 cells. The virtual reporter cassette tbox_c1 was constructed to match the original MPRA reporter architecture (Supplementary Figure 1). Each enhancer fragment was inserted upstream of a minimal promoter within the cassette, consistent with the original MPRA design, thereby reconstructing a complete promoter configuration for AlphaGenome inference in K562 cells. The VirDLA activity score was calculated from the predicted reporter signals and compared with linearized MPRA activity scores using Pearson and Spearman correlation analyses.

### 2.2 Genetic Algorithm–Driven Promoter Evolution

#### 2.2.1 Initial Population Construction

The initial promoter sequence was used as the seed sequence for population generation. The seed sequence itself was retained as the first individual in the population. The remaining N−1 individuals were generated by introducing 1–3 random point mutations into the seed sequence. For each mutation event, a nucleotide position was selected uniformly at random and replaced with A, C, G, or T, including the possibility of retaining the original nucleotide to simulate neutral mutations.

#### 2.2.2 Fitness Function Design

To preserve tissue specificity during evolution, candidate promoters were evaluated using a fitness function derived from VirDLA activity scores. Promoter selectivity was defined as:

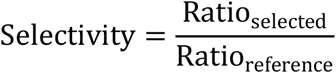

Where Ratio_selected_ and Ratio_reference_ denote the normalized VirDLA activity scores in the selected and reference tissues, respectively.

The overall fitness score was defined as:

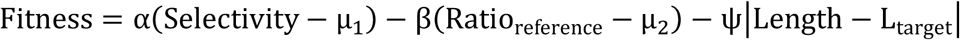

Where μ1 and μ2 are regularization terms introduced to prevent artificial inflation of selectivity caused by near-zero Ratioreference values. The coefficients α, β and ψ control the contributions of selectivity gain, off-target expression penalty, and length deviation penalty, respectively, while Ltarget denotes the target sequence length.

#### 2.2.3 Population Evaluation and Elite Preservation

At the beginning of each generation, all individuals in the population were evaluated using the fitness function described above. Fitness scores were calculated for every promoter sequence and stored for subsequent selection.

To prevent the loss of high-performing solutions, an elite preservation strategy was employed. Individuals were ranked in descending order according to fitness, and the top N_elite_ individuals (10% of the population size N ) were designated as elites. Elite individuals were temporarily removed from the evolutionary process and directly copied into the next generation after offspring production.

#### 2.2.4 Tournament Selection

Tournament selection was used to construct the mating pool. For each selection event, k individuals were sampled uniformly at random from the current population without replacement within the tournament, and the individual with the highest fitness score was selected. This procedure was repeated N times, where N denotes the population size, to generate a mating pool of equal size. Selection events were independent, allowing the same individual to be selected multiple times across different tournaments.

#### 2.2.5 Single-Point Crossover

Following tournament selection, offspring were generated using single-point crossover. Parent sequences were paired within the mating pool and recombined with probability Pc. If crossover was not performed, parental sequences were propagated unchanged to the offspring population. Population size was maintained throughout the evolutionary process.

#### 2.2.6 Mutation

Mutation was applied to offspring sequences following crossover. First, point mutations were introduced by traversing each nucleotide position independently and replacing the nucleotide with probability P ^m^ . This operation modified sequence composition without altering sequence length.

In addition to point mutations, structural mutation events were introduced with probability P ^i^ . For each structural mutation event, one of four mutation operators was selected at random: insertion, deletion, duplication, or recombination deletion.Insertion mutations introduced a random nucleotide fragment at a random position. Deletion mutations removed a contiguous sequence fragment. Duplication mutations copied a contiguous fragment and inserted the duplicated sequence into a nearby position. Recombination deletion retained sequence on one side of a randomly selected position while removing the opposite side.

Together, these mutation operators enabled exploration of both local sequence variation and large-scale structural changes during promoter evolution.

#### 2.2.7 Elite Reinsertion and Generation Update

Following crossover and mutation, the Nelite elite individuals preserved from the previous generation were reintroduced into the offspring population. The remaining population members consisted of newly generated offspring sequences.

After elite reinsertion, one evolutionary generation was considered complete and the resulting population was used as the input for the next generation. Population size remained constant throughout the optimization process, while the highest-fitness solutions were protected from loss due to stochastic genetic operations.

#### 2.2.8 Evolution Decoupling Environment

To avoid an inherent bias toward sequence expansion, we did not directly use senescent versus proliferating IMR90 cells as the optimization objective. Long promoters naturally contain more regulatory elements, and genome foundation models may therefore reward sequence length by integrating both causal regulatory features and correlated sequence features ^15^. Under such conditions, evolutionary optimization can converge toward increasingly long sequences rather than compact functional architectures.

To decouple promoter compression from direct senescence optimization, K562 cells were selected as an alternative optimization context. We do not consider K562 a biological surrogate of cellular senescence. Instead, K562 serves as a deliberately mismatched optimization environment that remains sufficiently distinct from the IMR90 senescence model while still sharing a subset of regulatory programs associated with p16 activation. This cross-context optimization strategy discourages simple length-driven overfitting arising from the accumulation of redundant regulatory motifs and favors the emergence of compact regulatory cores that retain functionality across distinct biological contexts.

#### 2.2.9 Workflow Overview

A schematic overview of the VirEvo framework is presented in Figure 2. Briefly, the wild-type promoter sequence was used as the initial seed to generate a starting population through random mutagenesis. Candidate promoters were subsequently evaluated using the VirDLA system, and fitness scores were calculated according to promoter selectivity, off-target activity, and sequence length constraints.

**Figure 1.**
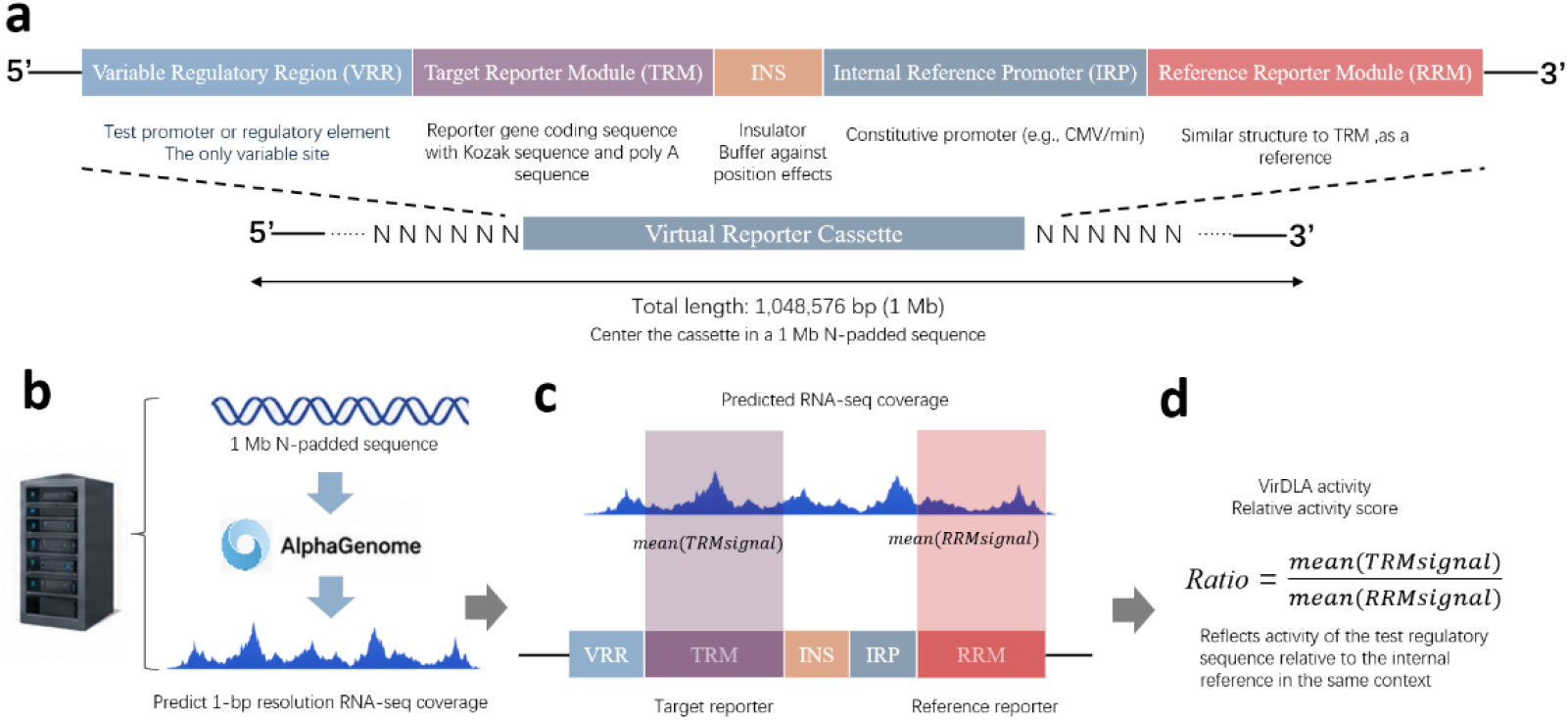
Overview of the Virtual Dual-Luciferase Assay (VirDLA). (a) Architecture of the virtual reporter cassette and construction of the AlphaGenome input sequence. The cassette is embedded at the center of a 1,048,576-bp N-padded sequence for prediction.(b) Prediction of strand-specific RNA-seq coverage across the virtual construct using AlphaGenome.(c) Extraction of mean predicted RNA-seq signals from the coding-sequence regions of the Target Reporter Module (TRM) and Reference Reporter Module (RRM).(d) Calculation of the VirDLA activity score as the ratio of the mean TRM signal to the mean RRM signal.

**Figure 2.**
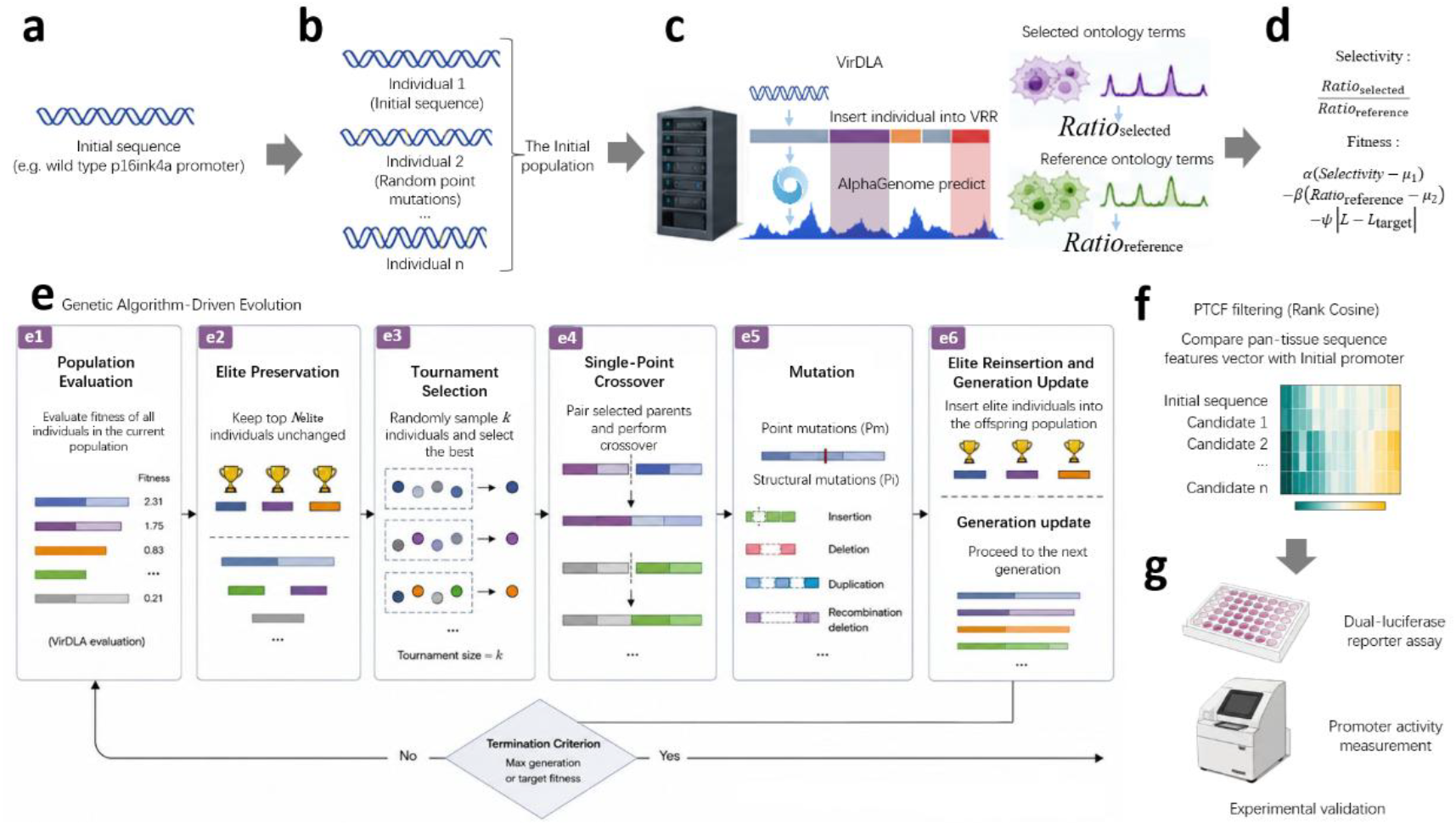
Schematic overview of the VirEvo framework. (a) Initial promoter sequence. (b) Initial population construction by random mutagenesis. (c) VirDLA-based activity prediction and selectivity evaluation. (d) Fitness calculation integrating selectivity, off-target activity, and sequence length. (e) Genetic algorithm–driven directed evolution. (f) PTCF screening using pan-tissue Rank Cosine similarity. (g) Experimental validation of selected promoter candidates by dual-luciferase reporter assay.

Evolutionary optimization was then performed through iterative cycles of tournament selection, crossover, mutation, and elite reinsertion. Following convergence, candidate promoters were subjected to PTCF screening based on pan-tissue Rank Cosine similarity. Selected candidates were selected for experimental validation using dual-luciferase reporter assays.

### 2.3 Pan-Tissue Consistency Filtering (PTCF)

Iterative optimization against predictive-model-derived fitness functions may introduce optimization bias and reward-hacking effects. To mitigate these risks, an orthogonal post-evolution screening strategy termed the Pan-Tissue Consistency Filter (PTCF) was developed. PTCF evaluates whether evolved promoters preserve the global tissue preference pattern of the wild-type promoter using pan-tissue VirDLA Sequence Feature Vectors.

#### 2.3.1 Construction of Pan-Tissue Sequence Feature Vectors

For the wild-type promoter, VirDLA activity scores were predicted across all available AlphaGenome tissue contexts and assembled into a pan-tissue Sequence Feature Vector:

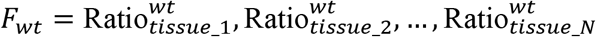

where 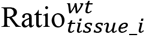 denotes the normalized VirDLA activity score of the wild-type promoter in tissue i, and N denotes the total number of evaluated tissue contexts.

#### 2.3.2 Candidate Profile Extraction

For all candidate promoters generated during evolutionary optimization (SRP1M– SRP4M and SRP6M–SRP8M), as well as the positive control CMV promoter and the negative-control random sequence (Ctrl), pan-tissue Sequence Feature Vectors were constructed using the same procedure described above.

For each sequence, VirDLA activity scores were predicted across all available AlphaGenome tissue contexts and assembled into a corresponding pan-tissue Sequence Feature Vector:

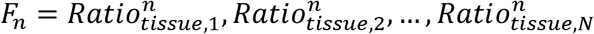

These vectors were subsequently used for Rank Cosine similarity analysis against the wild-type promoter profile.

#### 2.3.3 Rank Cosine Similarity Calculation

Because the objective of PTCF is to evaluate preservation of tissue-preference patterns rather than absolute promoter activity, direct cosine similarity on pan-tissue Sequence Feature Vectors may be confounded by differences in expression magnitude. Promoters exhibiting globally elevated activity can achieve high cosine similarity despite substantial changes in tissue-preference ordering. To reduce the influence of expression magnitude, each pan-tissue Sequence Feature Vector was transformed into a rank vector. Activity values were replaced by their ranks across all evaluated tissue contexts, where rank 1 corresponds to the lowest activity and rank N corresponds to the highest activity. Cosine similarity was then calculated using the resulting rank vectors:

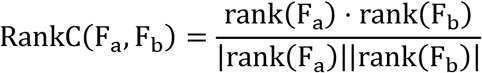

Rank Cosine similarity and Pearson correlation coefficients were calculated between each candidate promoter and the wild-type p16^INK4a^ promoter. The Rank Cosine value of the CMV promoter was used as the screening threshold. Candidate promoters exhibiting Rank Cosine values exceeding the CMV threshold and significant Pearson correlation (p < 0.05) were retained for subsequent experimental validation.

Because PTCF evaluates candidate promoters using pan-tissue Sequence Feature Vectors rather than the evolutionary fitness function, it provides an independent assessment of regulatory-profile conservation. By operating on rank-transformed Sequence Feature Vectors, Rank Cosine similarity reduces the influence of expression magnitude and emphasizes preservation of tissue-preference ordering.

### 2.4 Experimental Validation

The wild-type p16^INK4a^ promoter and the evolved SRP2M promoter were chemically synthesized and cloned into the pGL4.10[Fluc] luciferase reporter vector. Construct integrity was confirmed by restriction digestion and Sanger sequencing.

Reporter plasmids were transiently transfected into IMR90 human lung fibroblasts. Following transfection, cellular senescence was induced and verified by senescence-associated β-galactosidase (SA-β-gal) staining.

Promoter activities were quantified using dual-luciferase reporter assays in both proliferating and senescent cells. Relative promoter activity, basal leakage, induction activity, and senescence selectivity were subsequently calculated to evaluate the functional performance of SRP2M.

### 2.5 TFBS screening

TFBS analysis was performed using a custom motif scanning pipeline developed for this study. A library of 313 transcription factor binding motifs (Supplementary Table 3)was used for sequence scanning. Motif detection was based on fuzzy sequence matching, allowing a limited number of mismatches to accommodate biological sequence variability.

The wild-type p16^INK4a^ promoter, SRP2M promoter, and CMV promoter were analyzed using the same procedure. For each sequence, the total number of detected TFBSs was calculated and normalized by sequence length to obtain TFBS density.

TFBS counts, densities, and motif composition profiles were subsequently compared among promoters to characterize changes in regulatory architecture associated with promoter compression.

## RESULT

### 3.1 VirDLA benchmarking

To assess whether VirDLA captures experimentally measurable promoter activity, we benchmarked its predictions against an independent external dataset. The Sharpr-MPRA K562 scale-up dataset^14^ provides experimentally measured promoter activities for 1,000 human genomic fragments (295 bp each) in K562 cells. MPRA activity scores were linearized (2^x transformation) and used as ground truth.

Each promoter sequence was embedded into a virtual dual-luciferase reporter cassette, designated tbox_c1(Supplementary Figure 1), whose architecture mimics the vector backbone used in the original MPRA experiment(Supplementary Data 1). The complete cassette was N-padded to 1,048,576 bp and submitted to the AlphaGenome API with the ontology term EFO:0002067 (K562). The Fluc/Rluc ratio, computed as the mean RNA-seq signal over each CDS region, served as the VirDLA-predicted activity.

As expected, tbox_c1, which more closely recapitulates the original MPRA reporter architecture, achieved stronger agreement with experimentally measured activity than the alternative cassette.A parallel validation using an alternative cassette (tbox_h5) yielded consistent results.

VirDLA predictions showed significant correlation with experimentally measured MPRA activity (Figure 3). Pearson r = 0.531 (p = 5.7 × 10^-74^) and Spearman ρ = 0.481 (p = 4.1 × 10^-59^, n = 1,000), demonstrating that the virtual dual-luciferase framework captures both the linear trend and rank order of promoter activity in a zero-shot, fully external validation setting. The internal-reference normalization (Fluc/Rluc ratio) enables cross-sequence comparison that raw RNA-seq predictions alone cannot provide, and collectively demonstrate that VirDLA captures biologically meaningful variation in promoter activity despite relying exclusively on AlphaGenome outputs and without any task-specific model retraining.

**Figure 3.**
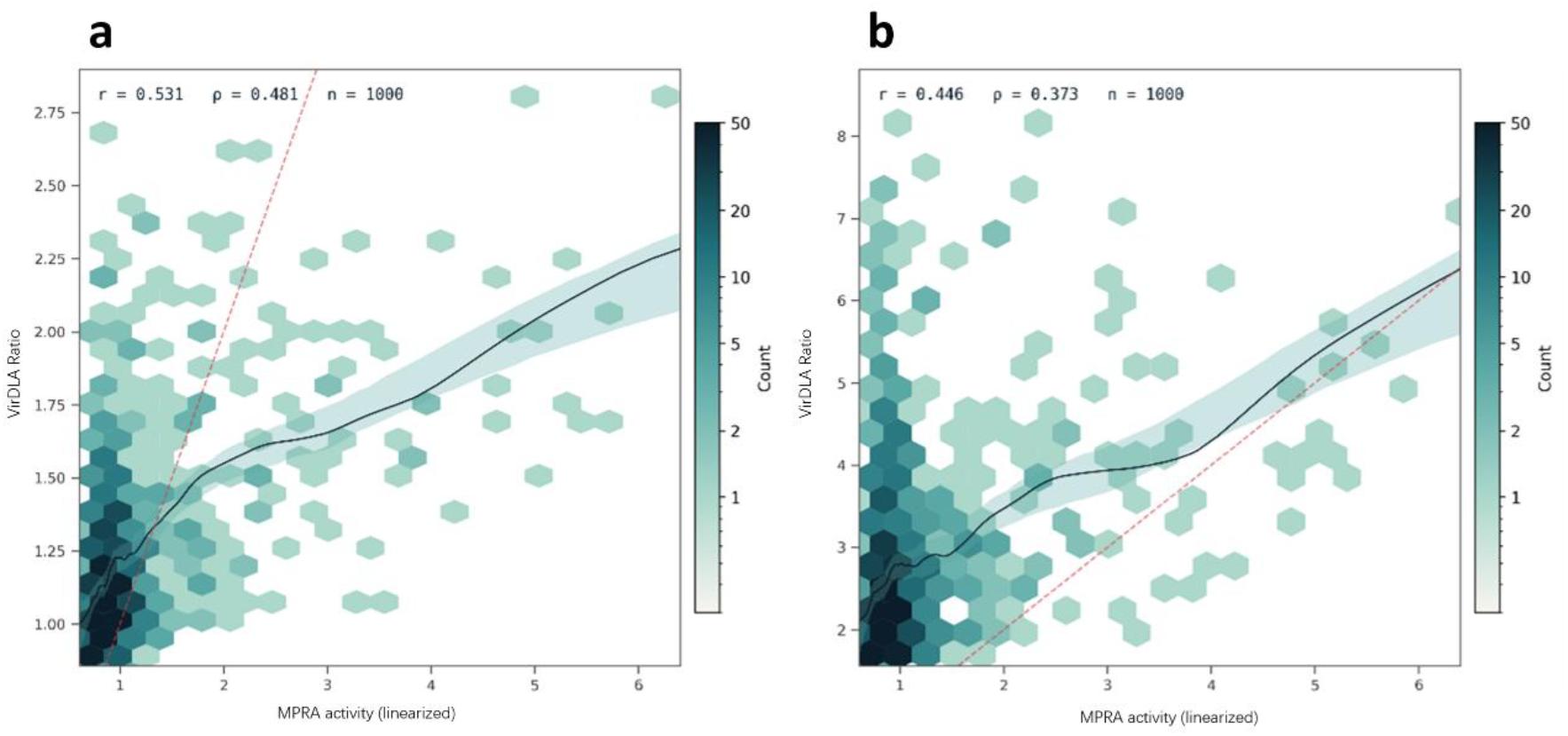
Zero-shot external validation of VirDLA using the SHARPR-MPRA K562 dataset (n = 1,000 independent 295-bp fragments). (a) Validation using tbox_c1, which employs the luciferase coding sequences from the MPRA vector backbone. Pearson r = 0.531 (p = 5.7 × 10^-74^), Spearman ρ = 0.481 (p = 4.1 × 10^-59^). (b) Validation using an alternative cassette, tbox_h5. Pearson r = 0.446 (p = 5.1 × 10^-50^), Spearman ρ = 0.373 (p = 2.0× 10^-34^). Teal curves show LOESS-smoothed trends with 95% confidence bands (bootstrap, 200 iterations); dashed red lines mark the diagonal y = x. Color intensity encodes local point density.

### 3.2 Virtual Evolution

Multiple evolutionary runs were performed using different optimization settings. Candidate promoters were periodically collected throughout the evolutionary process and subjected to PTCF screening. Across all runs, eight candidate promoters (SRP1M–SRP4M and SRP6M–SRP8M ; Supplementary Table 2) were recovered for downstream analysis. SRP5M was excluded because of an unintended sequence truncation introduced during sequence recovery.

### 3.3 PTCF Screening

PTCF screening was performed using Rank Cosine similarity across all recovered candidate promoters and reference controls (Supplementary Table 2). Candidate promoters were ranked according to Rank Cosine similarity using the CMV promoter (RankC = 0.915) as an empirical reference rather than a statistically optimal threshold.As shown in Figure 5, only SRP2M exceeded this threshold, achieving a Rank Cosine similarity of 0.939, whereas all remaining candidates, including SRP4M and the negative control, fell below the benchmark. To further examine the biological consistency of the selected candidate, tissue-level activity rankings were compared across 273 tissue types. Figure 4 shows that SRP2M preserved a tissue-ranking pattern broadly consistent with that of the wild-type p16^INK4a^ promoter. Based on these results, SRP2M was selected as the lead synthetic promoter for downstream experimental validation.

**Figure 4.**
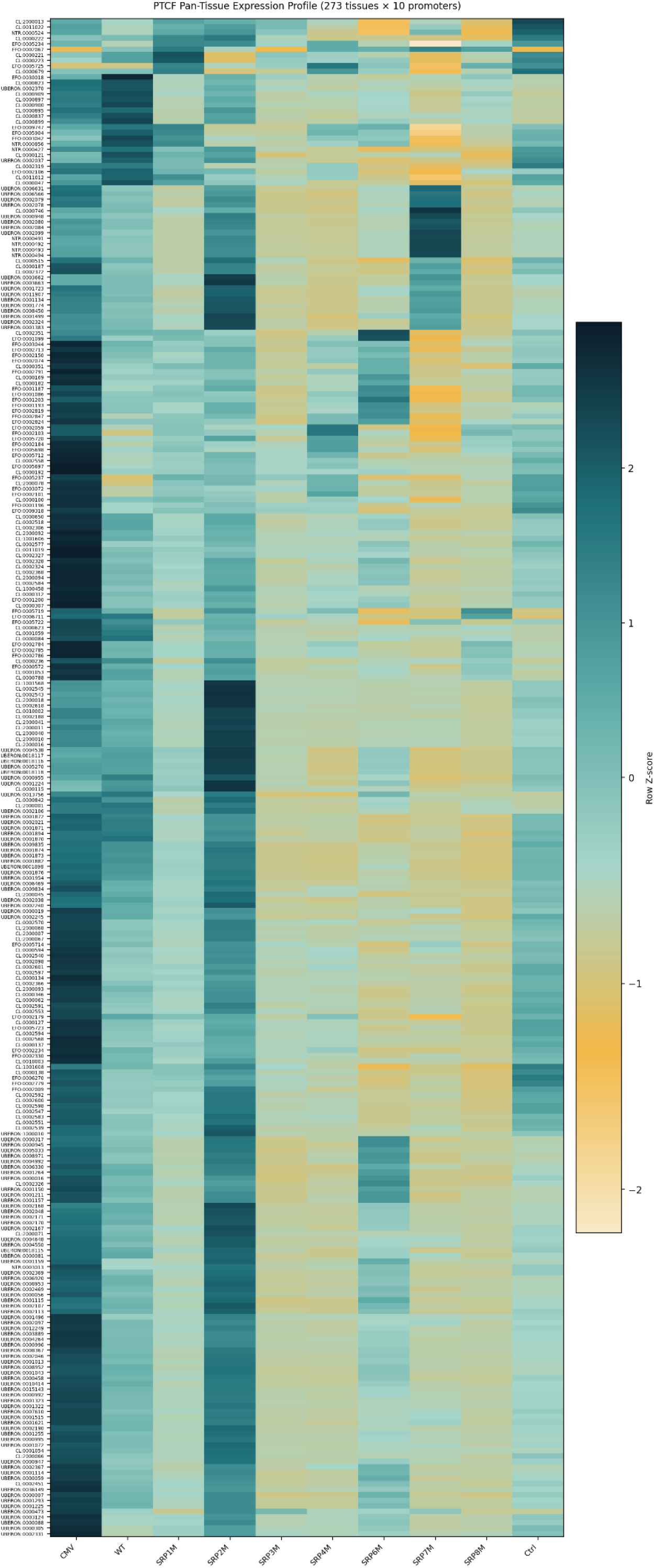
Pan-tissue expression profile heatmap of 10 promoters across 273 tissues. Row-wise Z-scores of Fluc/Rluc ratios. Columns are ordered as CMV, WT, SRP1M–SRP4M, SRP6M–SRP8M, and Ctrl. Rows are hierarchically clustered by expression pattern similarity.

**Figure 5.**
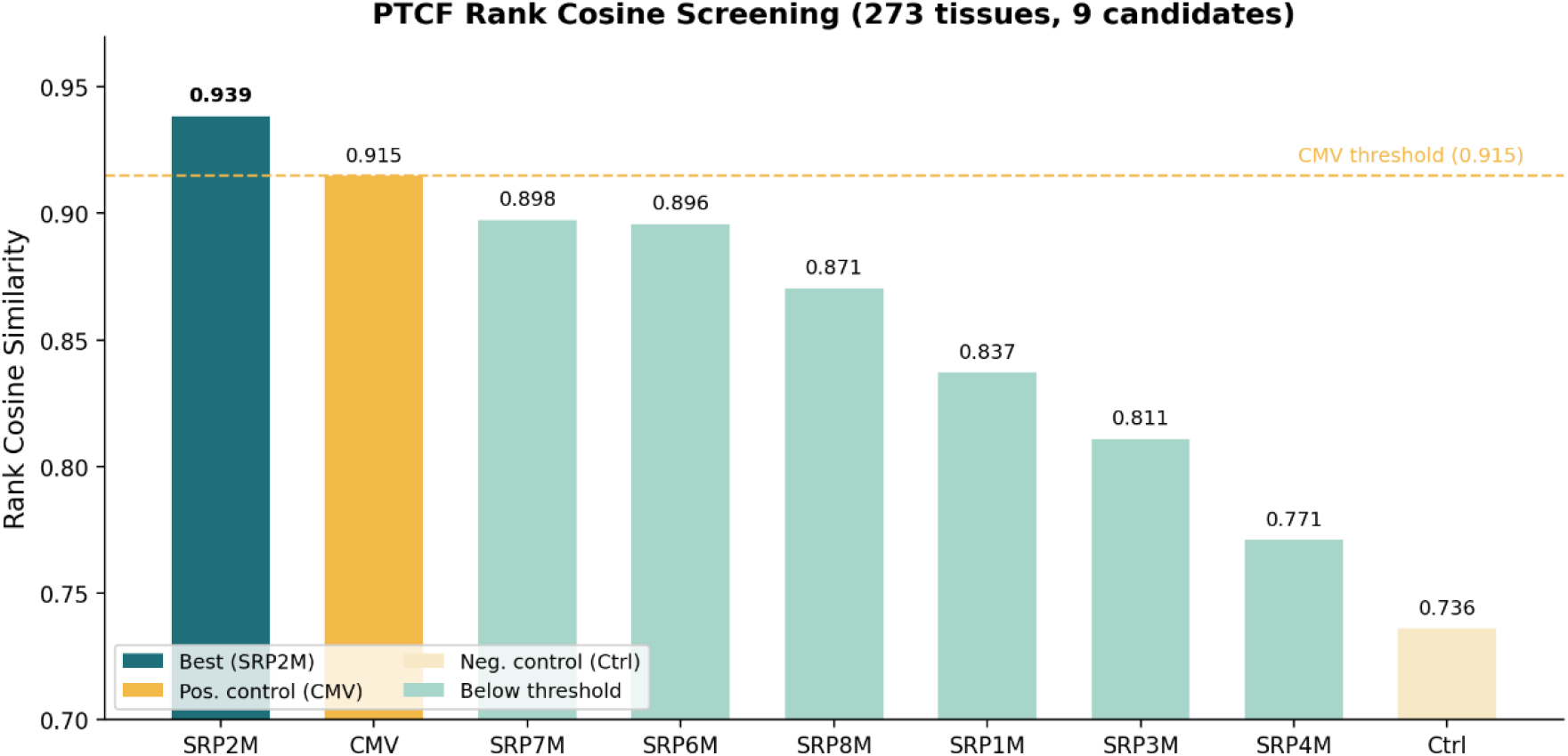
Rank Cosine similarity screening of all candidate promoters. Rank Cosine (RankC) values calculated relative to wild-type p16^INK4a^ promoter across 273 tissues.

### 3.4 Experimental validation

Promoter sequences were amplified by PCR and cloned into the pGL4.10[Fluc] dual-luciferase reporter vector using T4 DNA ligase. Recombinant plasmids were transformed into TOP10 competent cells, and positive clones were confirmed by colony PCR, restriction enzyme digestion, and Sanger sequencing.

To establish a senescence model for promoter validation, IMR90 human fibroblasts were treated with high-concentration D-galactose. Senescence induction was verified by senescence-associated β-galactosidase (SA-β-gal) staining. Control cells exhibited minimal SA-β-gal staining, whereas senescent cells displayed extensive blue precipitates and enlarged cellular morphology, consistent with canonical senescence phenotypes (Figure 6a,b). Quantitative analysis revealed that the percentage of SA-β-gal-positive cells increased from 14.16% in the control group to 87.95% in the senescent group (p < 0.01), confirming successful establishment of the senescence model (Figure 6c).

**Figure 6.**
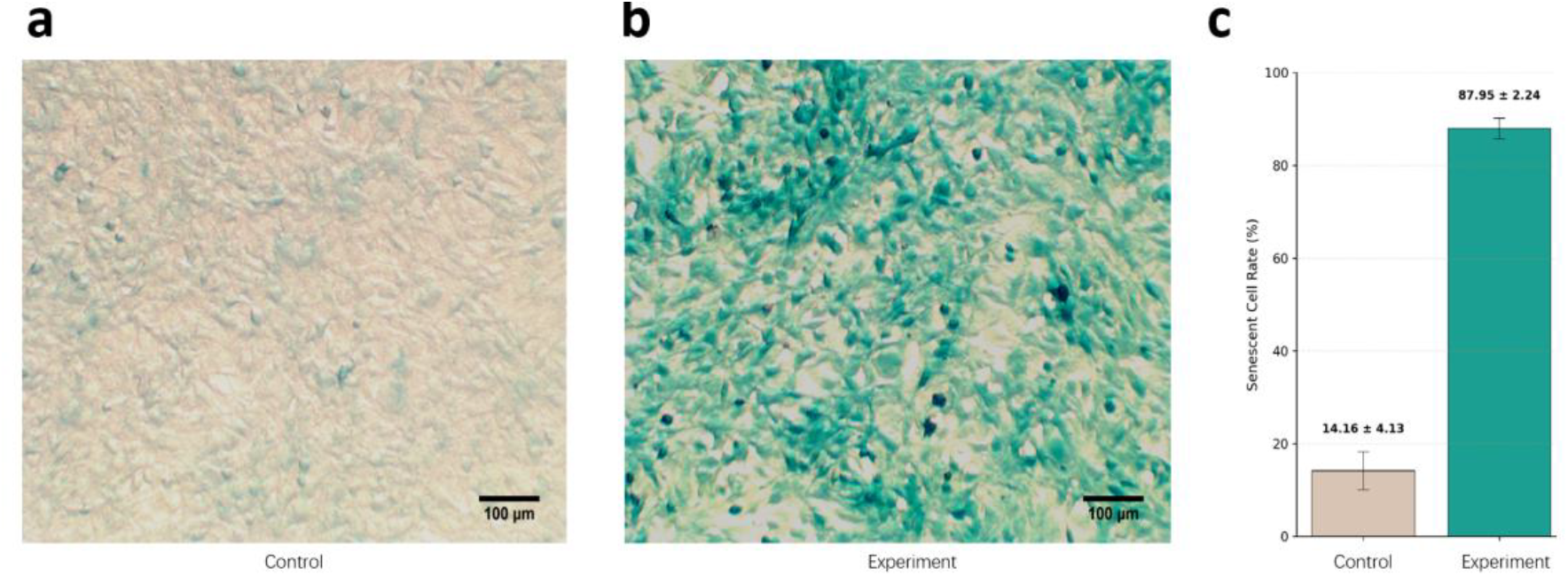
Experimental validation of plasmid construction and senescence induction. (a,b) Representative light microscopy images of SA-β-gal staining. Scale bars represent 100 μm. Figure a shows minimal blue precipitate (negative) in the control cells, while Figure b displays extensive dark blue precipitate (positive) in the senescent cells, indicating that the cells have entered a senescent state. (c) Quantification of the percentage of SA-β-gal-positive cells in the control group versus the senescent group. Data are presented as Mean ± SD, n=3.

The dual-luciferase reporter system was subsequently used to evaluate promoter activity in proliferating and senescent IMR90 cells (Figure 7). Compared with the wild-type p16^INK4a^ promoter, SRP2M retained approximately 77% of wild-type senescence selectivity while reducing promoter length to 14.1% of the original sequence. In addition, basal promoter activity in proliferating cells was reduced to approximately 52% of the wild-type level. These results demonstrate that substantial promoter compression can be achieved while preserving senescence-responsive transcriptional behavior.

**Figure 7.**
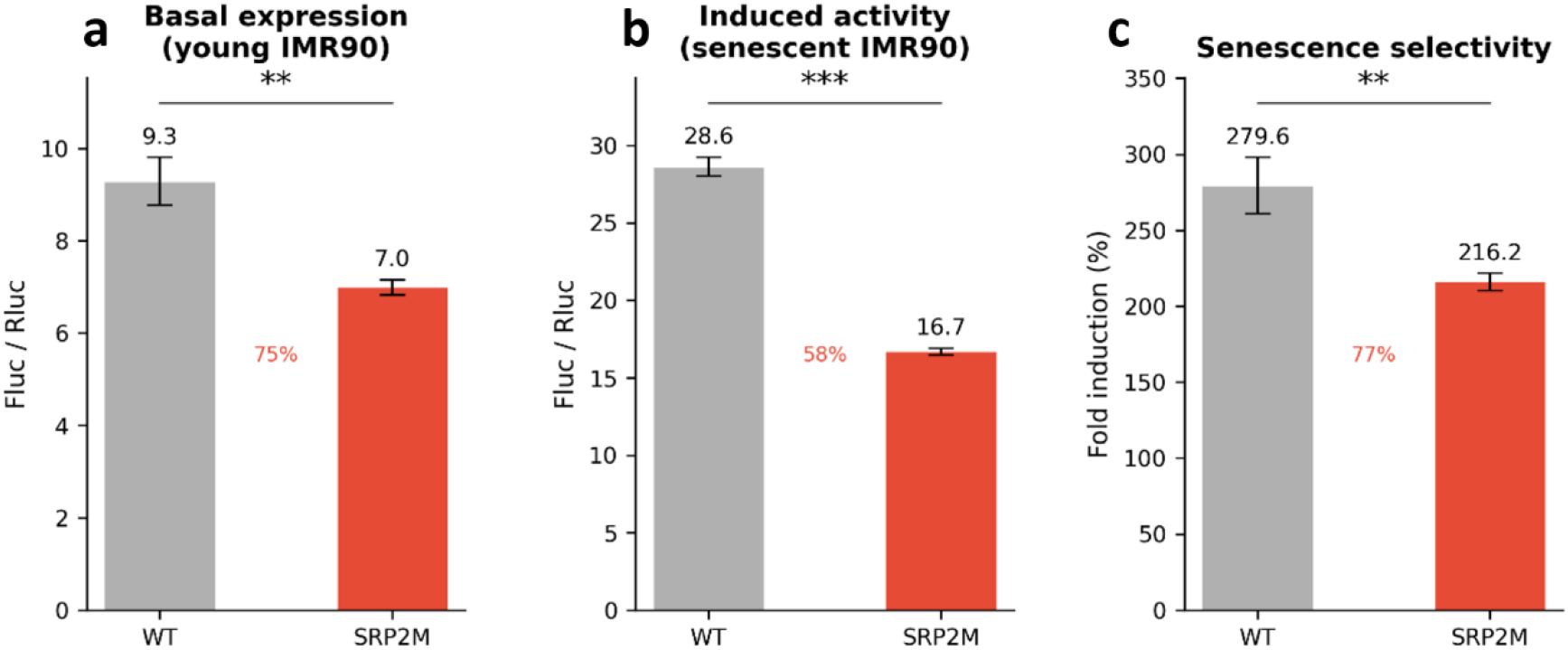
Basal expression, induced activity, and senescence selectivity of SRP2M. (a) Basal Fluc/Rluc activity in young IMR90 cells, reflecting promoter leakiness. (b) Induced Fluc/Rluc activity in senescent IMR90 cells. (c) Senescence selectivity index (fold induction) calculated as the ratio of induced to basal activity. All values are normalized to wild-type p16^INK4a^ (set to 100%). SRP2M achieves 77% of wild-type selectivity at 14% of the wild-type sequence length, with basal leakage reduced to 52% of the wild-type level.

### 3.5 TFBS screening

Based on a library of 313 transcription factor binding motifs (Supplementary Table 3), we compared the TFBS architectures of the wild-type p16^INK4a^ promoter (WT), SRP2M, and the CMV promoter. A total of 137 motif families (765 motif occurrences) were detected in WT, compared with 73 motif families (160 occurrences) in SRP2M and 82 motif families (277 occurrences) in CMV. Despite a reduction to only 14.1% of the wild-type sequence length, SRP2M retained 62 of 137 WT motif families (45.3%) and exhibited an approximately 3.8-fold increase in TFBS density relative to WT (Figure 8a,d).

**Figure 8.**
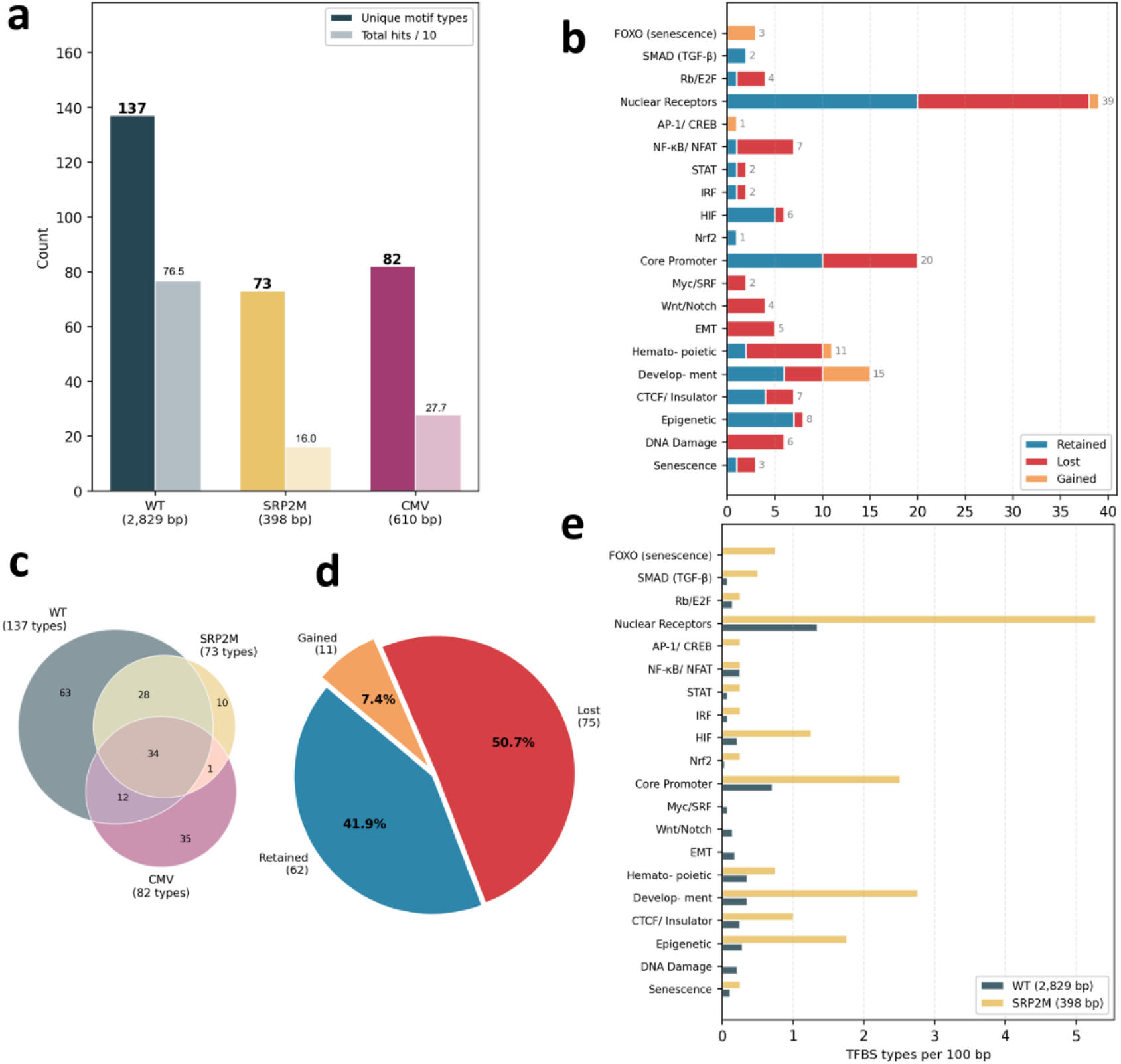
Comparative transcription factor binding site (TFBS) architecture analysis of the wild-type p16^INK4a^ promoter and the evolved SRP2M promoter. (a) Comparison of TFBS diversity and total TFBS occurrences detected in WT, SRP2M, and CMV promoters. (b) Retention, loss, and gain of TFBS families grouped by functional category. (c) Venn diagram showing overlap and uniqueness of TFBS repertoires among WT, SRP2M, and CMV promoters. (d) Distribution of TFBS fate following evolutionary compression, showing retained, lost, and newly gained motif families in SRP2M relative to the wild-type promoter. (e) Normalized TFBS family density (motif families per 100 bp) in WT and SRP2M promoters.

Analysis of motif-family composition revealed non-random patterns of TFBS retention and loss during evolutionary compression (Figure 8b,e). Several regulatory elements previously implicated in p16^INK4a^-associated pathways, including p14ARF-associated motifs, SMAD-family motifs, and E2F-related motifs, remained detectable in SRP2M. In contrast, numerous motifs associated with DNA-damage signaling, EMT-related transcription factors, and stress-responsive nuclear receptor subfamilies were absent from the compressed promoter. In addition, FOXO-family motifs (FOXO1/3/4), which were not detected in the wild-type promoter under the current scanning criteria, were identified in SRP2M.

Although TFBS scanning alone cannot establish causal regulatory mechanisms, these observations suggest that promoter compression did not occur through uniform motif loss. Instead, SRP2M retained a subset of the regulatory architecture present in the native promoter while simultaneously acquiring novel motif combinations. Together with the experimental observation that SRP2M preserved substantial senescence selectivity despite extensive sequence reduction, these results are consistent with the existence of a compact regulatory core within the p16^INK4a^ promoter.

## DISCUSSION

The development of compact and selective synthetic promoters remains a major challenge in gene therapy and synthetic biology. While recent advances in genome foundation models have substantially improved sequence-to-function prediction, these models are primarily designed for interpretation rather than engineering. In this study, we combined a virtual dual-luciferase assay framework (VirDLA), a genetic algorithm-based evolutionary engine (VirEvo), and an orthogonal pan-tissue screening strategy (PTCF) to establish a computational workflow for promoter compression and optimization. Using the human p16^INK4a^ promoter as a proof-of-concept system, we obtained SRP2M, a synthetic promoter with a length of only 398bp, representing 14.1% of the wild-type promoter length while retaining 77% of wild-type senescence selectivity.

Unlike conventional applications of AlphaGenome that rely directly on raw model outputs, VirDLA introduces an internal reference architecture inspired by dual-luciferase reporter assays. This design converts context-dependent prediction signals into relative activity measurements that can be compared across tissues and cellular states. Benchmarking against an independent SHARPR-MPRA dataset demonstrated a moderate positive correlation between VirDLA predictions and experimentally measured regulatory activities (Pearson r = 0.531, n = 1000 ). This level of agreement is expected because SHARPR-MPRA and VirDLA quantify regulatory activity using different experimental paradigms and different activity scales rather than identical measurements. Nevertheless, the observed positive correlation suggests that VirDLA captures biologically meaningful aspects of promoter function despite these methodological differences. Furthermore, evolutionary optimization depends primarily on preserving the relative ranking of candidate fitness rather than achieving perfectly calibrated quantitative predictions.

Optimization of a learned fitness proxy inevitably introduces the possibility of reward hacking, whereby evolved sequences exploit model-specific biases without preserving the intended biological function. To mitigate this risk, we introduced PTCF as an orthogonal validation layer independent of the VirEvo fitness function. Rather than focusing on absolute activity values, PTCF evaluates conservation of tissue-level activity rankings across 273 tissues and cell types. We selected Rank Cosine similarity because conventional similarity metrics are strongly influenced by expression magnitude, potentially favoring broadly active promoters over promoters that genuinely preserve tissue-specific regulatory patterns. By emphasizing relative ranking structure, PTCF provides an alternative criterion for assessing functional conservation and reduces the likelihood of selecting candidates that merely exploit properties of the prediction model.

We do not claim that genetic algorithms are the globally optimal search strategy for promoter engineering. Rather than benchmarking optimization algorithms, the objective of this study was to establish whether genome-foundation-model-guided evolutionary search can produce experimentally validated compact promoters.Genetic algorithms were chosen because promoter compression requires simultaneous exploration of point mutations and large structural alterations, including insertion, deletion, duplication, and truncation events. These operations naturally map onto evolutionary operators and allow sequence length to vary throughout optimization. The successful identification of SRP2M demonstrates the feasibility of this strategy, while systematic comparison against Bayesian optimization, simulated annealing, evolutionary strategies, or reinforcement learning remains an important direction for future work.

SRP2M also provides insight into promoter compressibility. Despite an overall sequence reduction of approximately 85.9%, SRP2M retained substantial functional characteristics of the wild-type promoter. TFBS analysis revealed that many senescence-associated regulatory elements were preserved or enriched, whereas numerous motifs associated with DNA damage response, stress signaling, and unrelated regulatory programs were lost. Although these observations do not establish causal mechanisms, they suggest that promoter function may be maintained through a condensed subset of regulatory interactions rather than through preservation of the complete native architecture. We therefore speculate that the combined pressures of sequence compression and cross-context optimization favored the retention of high-value regulatory elements while eliminating redundant regulatory information, ultimately yielding a compact regulatory core capable of preserving substantial promoter function.

Several limitations should be acknowledged. First, VirDLA benchmarking was performed using a single external dataset, and additional validation against diverse promoter activity datasets will be necessary to fully characterize predictive performance. We do not consider K562 a biological surrogate of senescence. Rather, K562 was used as a deliberately mismatched optimization environment that shares a subset of regulatory programs with p16 activation while remaining sufficiently distinct from IMR90 senescence. This decoupling discourages simple length-driven overfitting and favors the discovery of compact regulatory cores that generalize across contexts.Although this strategy successfully yielded a compact promoter with experimentally validated senescence responsiveness, future studies may benefit from incorporating more physiologically relevant senescence models directly into the optimization process. Third, the TFBS analyses are computational and do not establish causal regulatory mechanisms.

A central challenge in computational promoter engineering is not sequence generation itself, but candidate prioritization. Because the theoretical promoter search space is astronomically large whereas experimental validation is typically limited to a small number of constructs, computational frameworks that efficiently prioritize candidates are of substantial practical value. More broadly, this work suggests that genome foundation models may serve not only as tools for biological interpretation, but also as engines for biological design. Because VirEvo leverages pretrained genome foundation models rather than task-specific generative models, it can be readily adapted to different promoter engineering tasks without collecting task-specific training datasets or retraining models. As increasingly accurate genome foundation models become available, such computational design frameworks may accelerate the engineering of regulatory elements and broaden the application of AI-guided synthetic biology.

## ASSOCIATED CONTENT

### Supporting Information

Supplementary Table 1: Genetic algorithm parameters ; Supplementary Table 2 : Nucleotide sequences of all candidate and control promoters ; Supplementary Table 3 : Summary of the 313-motif TFBS library ; Supplementary Figure 1: Detailed schematic and annotations of the VirDLA reporter cassettes ; Supplementary Data 1: Complete benchmarking datasets comparing VirDLA predictions with MPRA measurements for tbox_c1 and tbox_h5 in TSV format ; Supplementary Data 2: Pan-tissue Activity Feature Vectors and expression matrices across 273 tissues and cell types in TSV format .

## AUTHOR INFORMATION

## NOTES

The author declares no competing financial interests.

