## Supporting Information for "Directed evolution of compact synthetic promoters via AlphaGenome and genetic algorithms"

Linxiao Nie[1,*]

[1]International College, Northeast Agricultural University, Harbin 150030, China.

### 1 Supplementary Table 1: Genetic algorithm parameters

| **Parameter** | **Value** |
|---|---|
| Population size ($N$) | 15 |
| Number of generations ($G$) | 50 |
| Crossover rate ($P_c$) | 0.6 |
| Point mutation rate ($P_m$) | 0.1 |
| Structural mutation rate ($P_i$) | 0.6 |
| Max structural mutation length | 50 bp |
| $\alpha$ (selectivity weight) | 1.0 |
| $\beta$ (leakiness weight) | 1.0 |
| $\psi$ (length penalty coefficient) | 0.2 |
| Target length ($L_{\text{target}}$) | 400 bp |
| Target selectivity | 0.5 |
| Target leakiness | 0.2 |

#### 2 Supplementary Table 2: Nucleotide sequences of all candidate and control promoters

| Name | Length | Sequence (5→3) |
|---|---|---|
| WT | 2829 | CGATGGCTCATGCCTGTAATCCCAGCACTTTGGGAGGCTAAGGTGGGTGGATCACCTGAGGTCAGGAGTTCG |
| | | AGACAAGCCTAGCCAACATAGTGAAACCCCGTCTCTACTAATAATACAAAAATTAGCTGGGTATGGCAGCAT |
| | | GTGCCTGTAATCCCAGCTACTCGGGAGGCTGAGGCAGGAGAATTGCTCGAACCCGGGAGGCGGAGGTTGCAG |
| | | TGAACCGAGAGAGATCGTGCGGTGCCATTTCACTCCAGCCTGGGCAACAGAGCGAAACTCCATCTCAAAAAA |
| | | ACACACAAAACAAACAAACAAAAAGAAAGAACCATTGTATTAGTGATGGAAATGTGTTCCCTCCCTCCCATC |
| | | CTGGCAACCACTTTCTTCCTCCTCCATCATAAAATATCTTAAACTAAACTAAAATAATTTTATTTATCGATA |
| | | GTTTGAATTTTCCCTATCATTGCTACACAGCTAATTGAGAGGTACCCCGAGGAAAATATAAATGGTACAGTA |
| | | ATGCATTGTAGATTTTAATAACATACTTGACATCCCAAATTGTTTTCATTGGCTTCATTTTAAAAACTACAT |
| | | GTTTTAAAATCAAGCAGACACTAAAAGTACAAGATATACTGGGTCTACAAGGTTTAAGTCAACCAGGGATTG |
| | | AAATATAACTTTTAAACAGAGCTGGATTATCCAGTAGGCAGATTAAGCATGTGCTTAAGGCATCAGCAAAGT |
| | | CTGAGCAATCCATTTTTTAAAACGTAGTACATGTTTTTGATAAGCTTAAAAAGTAGTAGTCACAGGAAAAAT |
| | | TAGAACTTTTACCTCCTTGCGCTTGTTATACTCTTTAGTGCTGTTTAACTTTTCTTTGTAAGTGAGGGTGGT |
| | | GGAGGGTGCCCATAATCTTTTCAGGGAGTAAGTTCTTCTTGGTCTTTCTTTCTTTCTTTCTTTCTTTTTTTC |
| | | TTGAGACCAAGTTTCGCTCTTGTCTCCCAGGCTGGAGTGCAATGGCGCGATCTCGGCTCACTGCAACCTCCG |
| | | CCTTCTCCTGGGTTCAAGCGATTCTCCTACATCAGCCTCCGAGTAGCTGGGATTACAGGCATGCGCCACCAA |
| | | GCCCCGCTAATTTTGTATTTTTTAGTAGAGACAGGGTTTCGCCATGTTGGTCAGGCTTGTCTCGAACTCCTG |
| | | GCCTCAGGTGATCCGCCTGTCTCGGCCTCCCAGAATGCTGGGATTATAGACGTGAGCCACCGCATCCGGACT |
| | | TTCCTTTTATGTAATAGTGATAATTCTATCCAAAGCATTTTTTTTTTTTTTTTTGAGTCGGAGTCTCATTCT |
| | | GTCACCCAGGCTGGAGGGTGGTGGCGCGATCTCGGCTTACTGCAACCTCTGCCTCCCGGGTTCAAGCGATTC |
| | | TCCTGCCTCAGCCTCCTGAGTAGCTGGAATTACACACGTGCGCCACCATGGCCAGCTAATTTTTGTATTTTT |
| | | AGTAGAGACGGGGTGTCACCATTTTGGCCAAGCTGGCCTCGAACTCCTGACCTCAGGTGATCTGCCCGCCTC |
| | | GGCTTCCCAAAGTGCTGGGATTACAGGTGTGAGCCACCGCGTCCTGCTCCAAAGCATTTTCTTTCTATGCCT |
| | | CAAAACAAGATTGCAAGCCAGTCCTCAAAGCGGATAATTCAAGAGCTAACAGGTATTAGCTTAGGATGTGTG |
| | | GCACTGTTCTTAAGGCTTATATGTATTAATACATCATTTAAACTCACAACAACCCCTATAAAGCAGGGGGCA |
| | | CTCATATTCCCTTCCCCCTTTATAATTACGAAAAATGCAAGGTATTTTCAGTAGGAAAGAGAAATGTGAGAA |
| | | GTGTGAAGGAGACAGGACAGTATTTGAAGCTGGTCTTTGGATCACTGTGCAACTCTGCTTCTAGAACACTGA |
| | | GCACTTTTTCTGGTCTAGGAATTATGACTTTGAGAATGGAGTCCGTCCTTCCAATGACTCCCTCCCCATTTT |
| | | CCTATCTGCCTACAGGCAGAATTCTCCCCCGTCCGTATTAAATAAACCTCATCTTTTCAGAGTCTGCTCTTA |
| | | TACCAGGCAATGTACACGTCTGAGAAACCCTTGCCCCAGACAGCCGTTTTACACGCAGGAGGGGAAGGGGAG |
| | | GGGAAGGAGAGAGCAGTCCGACTCTCCAAAAGGAATCCTTTGAACTAGGGTTTCTGACTTAGTGAACCCCGC |
| | | GCTCCTGAAAATCAAGGGTTGAGGGGGTAGGGGGACACTTTCTAGTCGTACAGGTGATTTCGATTCTCGGTG |
| | | GGGCTCTCACAACTAGGAAAGAATAGTTTTGCTTTTTCTTATGATTAAAAGAAGAAGCCATACTTTCCCTAT |
| | | GACACCAAACACCCCGATTCAATTTGGCAGTTAGGAAGGTTGTATCGCGGAGGAAGGAAACGGGGCGGGGGC |
| | | GGATTTCTTTTTAACAGAGTGAACGCACTCAAACACGCCTTTGCTGGCAGGCGGGGGAGCGCGGCTGGGAGC |
| | | AGGGAGGCCGGAGGGCGGTGTGGGGGGCAGGTGGGGAGGAGCCCAGTCCTCCTTCCTTGCCAACGCTGGCTC |
| | | TGGCGAGGGCTGCTTCCGGCTGGTGCCCCCGGGGGAGACCCAACCTGGGGCGACTTCAGGGGTGCCACATTC |
| | | GCTAAGTGCTCGGAGTTAATAGCACCTCCTCCGAGCACTCGCTCACGGCGTCCCCTTGCCTGGAAAGATACC |
| | | GCGGTCCCTCCAGAGGATTTGAGGGACAGGGTCGGAGGGGGCTCTTCCGCCAGCACCGGAGGAAGAAAGAGG |
| | | AGGGGCTGGCTGGTCACCAGAGGGTGGGGCGGACCGCGTGCGCTCGGCGGCTGCGGAGAGGGGGAGAGCAGG |
| | | CAGCGGGCGGCGGGGAGCAGC |
| SRP1M | 398 | AAAATACTATATTGTCAGAAATCAGGTCGTCTTGGACGGGATCTAGTCCCAGTTAAAAGGATTATCTTCCTT |
| | | CTCCACCTGCACGGGGGGTGTGAGTCACGGTACAGAACTTATCTCGTTTAACCCACACCTTCGAGCCGGAGG |
| | | ATTGGATTAAAGCCGTTAGTCACAAGCTTGCGCCAGAATTTGTTCGATGCGGGCGAAGTCAGGGCCCTCCTT |
| | | CGAGCTTGAACCCACAACTGCCTGGGATATGTTCGGAGTTAGGTTACGCCTCCACCATTCTCACGGTAAGTG |
| | | TTATGGTAATCTTATACGCTTAAGAGCGAGTATTTTTGGACGCAAGAATGGTAATCAGTAGGGGGTTTACCA |
| | | AGGTGAAAATTGGGAAATGGAAGGACGTCGGCACGAAA |

*Continued*

| Name | Length | Sequence |
|---|---|---|
| SRP2M | 398 | CCCCGAGGTCATTAGGAAGCCTCTTCTGGGTCTAAAGCAACGGAGCGGCTGTGAATGTCCGCAGCGCACAGA |
| | | GGGAAGGGCCGTCGCCAATTAGCTCTGAGGAAACTCCTTGTGATGGGGGTTGGTTCAATGAGGAAGGAAACG |
| | | GGCGGGACGCGGATTTATTTTTAACTTTAGCTAGAGGGTGGATATGTTTACCCTGTGGTACGCACGGAGTCC |
| | | ACCCTTGTGACTCGCAGGCCACATAAACGGCCCGCTAAGATCCTATGCCGCGGTCCTACTACCCCATGCTGG |
| | | TAAGATCCAACTGGGCGTGCGACTAGGAAATATGAAATAGCATGATCCCGACTGTTATCTCCTTCAAAGCAG |
| | | GGTCGTAGACGCAAGAGATCAGACAGCTAGAATTATGG |
| SRP3M | 401 | TACTCCCTTGTCCGCCTTAGCCCAGGGCGAATAAGCATCCGCAAAACACCTCAAGGCATGCTGCAGCGATCG |
| | | TGTTACTGATCTTGCTGAAGACGTTACGACTGGCCCATCGGAGATTATCCAAAGCATTTCAACTCTCTGAAA |
| | | TGCATGAAGTTAAATTTCAGCACGATTCAGGGACTTTAACAACCCAACCTTAACAACAGATAGTGGCGGGCT |
| | | CAGGGATCGGTGCTAACACTGCCGTGGGTTGAGTAGATGGTACGAATATTTAGTTAAAGATCCTTGGACCAA |
| | | CTAGCGGATTTGCGAGTTTAGTCCCAGTTTGAGCCTTCTTACGGTTATACACATCCGTTAAAAGGGAAATCT |
| | | AAATCGGCTTGCTTCCGTTCGTTATAACGACACTGCCGGTA |
| SRP4M | 400 | CCTAGCGATGGAGGTACGTTATGTTTGGGAGTACCTGATCATCTCACTAGTCGCGGAGACGTTGCATGTACC |
| | | GTTGTGGGGGAAAACTCGCACCAAAAGATGCCTTATTAAGAGAAGCCGACTCCACGAACCACACCCGGAAGC |
| | | TCATTAGCATATTTATCTAGAGGATCCCCGTTGATTAGTCATGTGGTTCCTCAAACGGTTGTAAAGCACCTG |
| | | GCTTGAGTGCTGCAGTGTACAGCAGATTCAGCCGACACATTGCCTTCCGTCCCGAGACTAACGGGATGCGCA |
| | | CGTCCTTCCACAACATCCGGGGACTGGTGGGTACGACCTGTGCTCGGACCTCTTTAGCCCGCGATCGTCACT |
| | | CTCTCACCACGAGCGTGGACCGTTCGGAAATTCATAGGTA |
| SRP6M | 397 | AACATACTATATTGTCAAAAATCAAGTCGTCTAGGACAGGCTCTACGCCCAGCTAAAAGGGGTATCTTCCTT |
| | | TTACACCTGCACGGGGGGTGTGAGTCACAGTACAAGGCTTATCTCGTTTAACCCACACCTTCGATCCAGAGG |
| | | ATTCGAGTAAAGCCGTTAGTCACAAGCTTGCGCAAGAGGTTGCTCGATGCGGGTGACGTCAGGGCCCTCCTT |
| | | CGACCTTGAACACACACCTGCCCGGGAACTGTGCGGAGTTAGGTTACGCTCCGAACATTCTCACGGTAAGAG |
| | | TTAGGGAATTCTTATATGCTTAAGGGCGAGTATTTTTAGAAGCACCAATGGTCCTCAGTCGAGAGTCACCTG |
| | | GCCGCAAATTGGAAGATGGAAGGACGTCGGCAAGAAA |
| SRP7M | 400 | CATGTAGATACCTGGGCGCCACTAGTGTCAGTGCGCAATTATCCAATCAATTTAGCAAATCTGTAAGGGTAA |
| | | CTTAACACGTACTAGGTGTAAGTCCCCGGAACGGGCTGATGTGTGTTTATCTCAATCATAGCATGGCAATGG |
| | | ACTGCATAAAAGAGCACTTGGGTTGCCCTTAGTGACATAAAAAGCTCACCCTTAACTAAAAGTTCCTGCGCC |
| | | TTTAGCGTTCGACCGCCGCGTGTCATCGGGCGTCTTTTTTTCTATAAATAGTGTCCATGCCTCTGGGACAGC |
| | | CGAGTGCAGCTGCTATTAAAGATAAGGTTTGTGCCTCGCAACGGGACCTGAAATTCGTTGGACACTTGAACC |
| | | TCTAGCGAATAATTTCCGTTAATCGTCACGAGCGACGATA |
| SRP8M | 400 | CGGCGAGATCGAGGTACAGTTTGTTTGAGTGTACATGATCAGCTGAATAGTCCCGGAGTCCTTGCATGTAGC |
| | | AGCACAAGGGAACACTCTCAACGAAAGAACCTTTAGCAAGAGAAGTAGATTACACGAACCGCATCCGGAACC |
| | | TCATTAGCATACTTATCTAGAGGCTCCCAATAGAGTATTCACGTAGTTCCTCAAACGGCTGTGAAGCACCTG |
| | | GCTTGAGTGCTGGAGTGTACAGCAGCTTCACCCAACATATCGCCTGCCCTCCCGCGACTTAAGAGCTGAGGA |
| | | CGTCCTTCCAAAACATTTCGGGACTGGTGGGTGCGACCCGTGTTTGGACCCCATTAGCCCGCGATACACACT |
| | | CTCTATCCACGATCCTAGACCGATCTGAAATTCATAGGTA |
| Contrl | 199 | NNNNNNNNNNNNNNNNNNNNNNNNNNNNNNNNNNNNNNNNNNNNNNNNNNNNNNNNNNNNNNNNNNNNNNNN |
| | | NNNNNNNNNNNNNNNNNNNNNNNNNNNNNNNNNNNNNNNNNNNNNNNNNNNNNNNNNNNNNNNNNNNNNNNN |
| | | NNNNNNNNNNNNNNNNNNNNNNNNNNNNNNNNNNNNNNNNNNNNNNNNNNNNNNN |
| CMV | 609 | CACCTATTGACTAGTTATTAATAGTAATCAATTACGGGGTCATTAGTTCATAGCCCATATATGGAGTTCCGC |
| | | GTTACATAACTTACGGTAAATGGCCCGCCTGGCTGACCGCCCAACGACCCCCGCCCATTGACGTCAATAATG |
| | | ACGTATGTTCCCATAGTAACGCCAATAGGGACTTTCCATTGACGTCAATGGGTGGAGTATTTACGGTAAACT |
| | | GCCCACTTGGCAGTACATCAAGTGTATCATATGCCAAGTACGCCCCCTATTGACGTCAATGACGGTAAATGG |
| | | CCCGCCTGGCATTATGCCCAGTACATGACCTTATGGGACTTTCCTACTTGGCAGTACATCTACGTATTAGTC |
| | | ATCGCTATTACCATGGTGATGCGGTTTTGGCAGTACATCAATGGGCGTGGATAGCGGTTTGACTCACGGGGA |
| | | TTTCCAAGTCTCCACCCCATTGACGTCAATGGGAGTTTGTTTTGGCACCAAAATCAACGGGACTTTCCAAAA |
| | | TGTCGTAACAACTCCGCCCCATTGACGCAAATGGGCGGTAGGCGTGTACGGTGGGAGGTCTATATAAGCAGA |
| | | GCTCGTTTAGTGAACCGTCAGATCGGCAATCCG |

### 3 Supplementary Table 3: Summary of the 313-motif TFBS library

| Category | Count | Example Motif(s) |
|---|---|---|
| Nuclear Receptors | 30 | RAR, RXR, PPAR, ER, GR, LXR, FXR, VDR, ROR, ERR, etc. |
| Epigenetic / Chromatin | 30 | EZH2, DNMT1/3, HDAC1/2, SIRT1/6, BRG1, MLL1/2, TET1/2, etc. |
| bHLH (Myc/HIF/TCF) | 29 | MYC, MAX, HIF1$\alpha$, TCF3/4/12, TWIST, HES, HEY, USF, SRE, SRF |
| bZIP (AP-1/CREB/ATF) | 26 | AP-1, CREB, ATF2/3/4, c-FOS, c-JUN, FRA1/2, BACH1/2, XBP1, NRF2 |
| Forkhead / HNF | 21 | HNF1$\alpha$, HNF4$\alpha$, FOXA1/2, FOXO1/3, FOXM1 |
| Core Promoter Elements | 18 | TATA, TBP, Inr, DPE, MTE, BRE, DCE, TCT, XCPE1 |
| Zinc Finger | 13 | SP1, KLF4/5, YY1, REST, ZIC1–5 |
| NF-$\kappa$B | 13 | NF$\kappa$B (p65, p50, c-Rel, RelB) |
| ETS | 13 | ETS1/2, ELK1, GABP$\alpha$, ERG, FLI1, PU.1, ESE1/2/3 |
| C/EBP | 10 | C/EBP$\alpha$, C/EBP$\beta$, C/EBP$\delta$, DBP |
| STAT | 9 | STAT1–6 |
| p53 Family | 9 | p53, p63, p73 |
| IRF / ISRE | 9 | IRF1–9, ISRE |
| Cell Cycle / DNA Damage | 9 | ATM, ATR, CHK1/2, $\gamma$H2AX, p21, p16, p14 |
| E2F / DP | 8 | E2F1–8, DP1 |
| Homeobox / POU | 8 | OCT4, NANOG, PAX5/6, HOXA9, HOXB4 |
| GATA | 6 | GATA1–6 |
| NFAT | 6 | NFATc1–c4, NFATC1/2 |
| SMAD (TGF-$\beta$) | 6 | SMAD1/5, SMAD2/3, SMAD4 |
| SOX / SRY | 5 | SOX2/9/10/17, SRY |
| CTCF / Cohesin | 4 | CTCF, CTCFL |
| EMT (SNAIL/ZEB) | 4 | SNAIL1, SLUG, ZEB1 |
| NFY (CCAAT-box) | 2 | NFY |
| RUNX | 2 | RUNX1, RUNX2 |
| Wnt (LEF/$\beta$-catenin) | 2 | LEF1, $\beta$-catenin/TCF |
| T-box | 2 | TBX5, BRACHYURY (T) |
| Notch (RBPJ) | 2 | RBPJ |
| SASP / Senescence | 2 | SASP-IL6, SASP-IL8 |
| Other | 15 | AR, BCL6, EBF1, Elk1, c-Fos, c-Jun, Fra1, Fra2, JunB, JunD, MEF2C, MEF2D, Myc |
| **Total** | **313** | |

### 4 Supplementary Figure 1: Detailed schematic and annotations of the VirDLA reporter cassettes

##### Modular architecture

The VirDLA reporter cassette consists of five functional modules arranged from upstream to downstream:

**VRR** (Variable Regulatory Region) → **TRM** (Target Reporter Module) → **6×cHS4 Insulator** → **IRP** (Internal Reference Promoter, CMV minimal) → **RRM** (Reference Reporter Module)

##### A. tbox_c1 (4,762 bp) — MPRA-mimicking architecture

Designed to match the original MPRA vector backbone. The minimal promoter is used to recapitulate the original MPRA experimental configuration.

**[VRR: enhancer] → [TRM: minP + Fluc + SV40pA] → [6×cHS4] → [IRP: CMV minimal] → [RRM: Rluc + hGHpA]**

**TRM (Fluc)** (1–1839, 1839 bp) — 64 bp minP + Fluc CDS + SV40pA

```
tagagggtatataatggaagctcgacttccagcttggcaatccggtactgttggtaaagccaccatggaagatgccaaaaacattaagaa
gggcccagcgccattctacccactcgaagacgggaccgccggcgagcagctgcacaaagccatgaagcgctacgccctggtgcccggcac
catcgcctttaccgacgcacatatcgaggtggacattacctacgccgagtacttcgagatgagcgttcggctggcagaagctatgaagcg
ctatgggctgaatacaaaccatcggatcgtggtgtgcagcgagaatagcttgcagttcttcatgcccgtgttgggtgccctgttcatcgg
tgtggctgtggccccagctaacgacatctacaacgagcgcgagctgctgaacagcatgggcatcagccagcccaccgtcgtattcgtgag
caagaaagggctgcaaaagatcctcaacgtgcaaaagaagctaccgatcatacaaaagatcatcatcatggatagcaagaccgactacca
gggcttccaaagcatgtacaccttcgtgacttcccatttgccacccggcttcaacgagtacgacttcgtgcccgagagcttcgaccggga
caaaaccatcgccctgatcatgaacagtagtggcagtaccggattgcccaagggcgtagccctaccgcaccgcaccgcttgtgtccgatt
cagtcatgcccgcgaccccatcttcggcaaccagatcatccccgacaccgctatcctcagcgtggtgccatttcaccacggcttcggcat
gttcaccacgctgggctacttgatctgcggctttcgggtcgtgctcatgtaccgcttcgaggaggagctattcttgcgcagcttgcaaga
ctataagattcaatctgccctgctggtgcccacactatttagcttcttcgctaagagcactctcatcgacaagtacgacctaagcaactt
gcacgagatcgccagcggcggggcgccgctcagcaaggaggtaggtgaggccgtggccaaacgcttccacctaccaggcatccgccaggg
ctacggcctgacagaaacaaccagcgccattctgatcacccccgaaggggacgacaagcctggcgcagtaggcaaggtggtgcccttctt
cgaggctaaggtggtggacttggacaccggtaagacactgggtgtgaaccagcgcggcgagctgtgcgtccgtggccccatgatcatgag
cggctacgttaacaaccccgaggctacaaacgctctcatcgacaaggacggctggctgcacagcggcgacatcgcctactgggacgagga
cgagcacttcttcatcgtggaccggctgaagagcctgatcaaatacaagggctaccaggtagccccagccgaactggagagcatcctgct
gcaacaccccaacatcttcgacgccggggtcgccggcctgcccgacgacgatgccggcgagctgcccgccgcagtcgtcgtgctggaaca
cggtaaaaccatgaccgagaaggagatcgtggactatgtggccagccaggttacaaccgccaagaagctgcgcggtggtgttgtgttcgt
ggacgaggtgcctaaaggactgaccggcaagttggacgcccgcaagatccgcgagattctcattaaggccaagaagggcggcaagatcgc
cgtgtaataagatacattgatgagtttggacaaaccacaactagaatgcagtgaaaaaaatgctttatttgtgaaatttgtgatgctatt
gctttatttgtaaccattataagctgcaataaacaagtt
```

**6×cHS4 Insulator** (1840–3273, 1434 bp)

```
gatccgtcgaccttttccccgtatcccccaggtgtctgcaggctcaaagagcagcgagaagcgttcagaggaaagcgatcccgtgccac
cttccccgtgcccgggctgtccccgcacgctgccggctcggggatgcgggggggagcgccggaccggagcggagccccgggcggctcgctg
ctgcccccctagcgggggagggacgtaattacatccctgggggctttgggggggggctgtccctggcaatcttttccccgtatcccccag
gtgtctgcaggctcaaagagcagcgagaagcgttcagaggaaagcgatcccgtgccaccttccccgtgcccgggctgtccccgcacgctg
ccggctcggggatgcgggggggagcgccggaccggagcggagccccgggcggctcgctgctgcccccctagcgggggagggacgtaattaca
tccctgggggctttgggggggggctgtccctcggttttccccgtatcccccaggtgtctgcaggctcaaagagcagcgagaagcgttca
```

gaggaaagcgatcccgtgccaccttccccgtgcccgggctgtccccgcacgctgccggctcggggatgcggggggagcgccggaccggag
cggagccccgggcggctcgctgctgccccctagcgggggagggacgtaattacatccctgggggctttggggggggctgtcccttactt
ttccccgtatccccccaggtgtctgcaggctcaaagagcagcgagaagcgttcagaggaaagcgatcccgtgccaccttccccgtgcccg
ggctgtccccgcacgctgccggctcggggatgcggggggagcgccggaccggagcggagccccgggcggctcgctgctgccccctagcgg
gggagggacgtaattacatccctgggggctttggggggggctgtcccttgtttttccccgtatccccccaggtgtctgcaggctcaaag
agcagcgagaagcgttcagaggaaagcgatcccgtgccaccttccccgtgcccgggctgtccccgcacgctgccggctcggggatgcggg
gggagcgccggaccggagcggagccccgggcggctcgctgctgccccctagcgggggagggacgtaattacatccctgggggctttgggg
ggggctgtcccttggttttccccgtatccccccaggtgtctgcaggctcaaagagcagcgagaagcgttcagaggaaagcgatcccgtg
ccaccttccccgtgcccgggctgtccccgcacgctgccggctcggggatgcggggggagcgccggaccggagcggagccccgggcggctc
gctgctgccccctagcgggggagggacgtaattacatccctgggggctttggggggggctgtcccttaaagccaccggcaatc

**IRP (CMV minimal)** (3274–3337, 64 bp)
tagagggtatataatggaagctcgacttccagcttggcaatccggtactgttggtaaagccacc

**RRM (Rluc)** (3338–4762, 1425 bp)
gcacctgcaggCatggcttccaaggtgtacgaccccgagcaacgcaaacgcatgatcactgggcctcagtggtgggctcgctgcaagcaa
atgaacgtgctggactccttcatcaactactatgattccgagaagcacgccgagaacgccgtgatttttctgcatggtaacgctacctcc
agctacctgtggaggcacgtcgtgcctcacatcgagcccgtggctagatgcatcatccctgatctgatcggaatgggtaagtccggcaag
agcgggaatggctcatatcgcctcctggatcactacaagtacctcaccgcttggttcgagctgctgaaccttccaaagaaaatcatcttt
gtgggccacgactgggggctgctctggcctttcactacgcctacgagcaccaagacaggatcaaggccatcgtccatatggagagtgtc
gtggacgtgatcgagtcctgggacgagtggcctgacatcgaggaggatatcgccctgatcaagagcgaagagggcgagaaaatggtgctt
gagaataacttcttcgtcgagaccgtgctcccaagcaagatcatgcggaaactggagcctgaggagttcgctgcctacctggagccattc
aaggagaagggcgaggttagacggcctaccctctcctggcctcgcgagatccctctcgttaagggaggcaagcccgacgtcgtccagatt
gtccgcaactacaacgcctaccttcgggccagcgacgatctgcctaagctgttcatcgagtccgaccctgggttcttttccaacgctatt
gtcgagggagctaagaagttccctaacaccgagttcgtgaaggtgaagggcctccacttcctccaggaggacgctccagatgaaatgggt
aagtacatcaagagcttcgtggagcgcgtgctgaagaacgagcagtaagggtggcatccctgtgacccctccccagtgcctctcctggcc
ctggaagttgccactccagtgcccaccagccttgtcctaataaaattaagttgcatcattttgtctgactaggtgtccttctataatatt
atggggtggaggggggtggtatggagcaaggggcaagttgggaagacaacctgtagggcctgcggggtctattgggaaccaagctggagt
gcagtggcacaatcttggctcactgcaatctccgcctcctgggttcaagcgattctcctgcctcagcctcccgagttgttgggattccag
gcatgcatgaccaggctcagctaatttttgtttttttggtagagacggggtttcaccatattggccaggctggtctccaactcctaatct
caggtgatctacccaccttggcctcccaaattgctgggattacaggcgtgaaccactgctcccttccctgtcctt

#### B. tbox_h5 (5,065 bp) — Complete humanized architecture

All non-coding regions humanized. The test promoter inserted at the VRR directly drives hluc2 — the TRM contains **no** embedded minimal promoter.

**[VRR: test promoter] → [TRM: hluc2 + hGHpA] → [6×cHS4] → [IRP: CMV minimal] → [RRM: rluc2 + hGHpA]**

**TRM (hluc2)** (1–2142, 2142 bp) — *no embedded promoter*
gcacctgcaggCatggaagatgccaaaaacattaagaagggcccagcgccattctacccactcgaagacgggaccgccggcgagcagctg
cacaaagccatgaagcgctacgccctggtgcccggcaccatcgcctttaccgacgcacatatcgaggtggacattacctacgccgagtac
ttcgagatgagcgttcggctggcagaagctatgaagcgctatgggctgaatacaaaccatcggatcgtggtgtgcagcgagaatagcttg
cagttcttcatgcccgtgttgggtgccctgttcatcggtgtggctgtggccccagctaacgacatctacaacgagcgcgagctgctgaac
agcatgggcatcagccagcccaccgtcgtattcgtgagcaagaaagggctgcaaaagatcctcaacgtgcaaaagaagctaccgatcata
caaaagatcatcatcatggatagcaagaccgactaccagggcttccaaagcatgtacaccttcgtgacttcccatttgccacccggcttc
aacgagtacgacttcgtgcccgagagcttcgaccgggacaaaaccatcgccctgatcatgaacagtagtggcagtaccggattgcccaag
ggcgtagccctaccgcaccgcaccgcttgtgtccgattcagtcatgcccgcgaccccatcttcggcaaccagatcatccccgacaccgct
atcctcagcgtggtgccatttcaccacggcttcggcatgttcaccacgctgggctacttgatctgcggctttcgggtcgtgctcatgtac
cgcttcgaggaggagctattcttgcgcagcttgcaagactataagattcaatctgccctgctggtgcccacactatttagcttcttcgct

aagagcactctcatcgacaagtacgacctaagcaacttgcacgagatcgccagcggcggggcgccgctcagcaaggaggtaggtgaggcc
gtggccaaacgcttccacctaccaggcatccgccagggctacggcctgacagaaacaaccagcgccattctgatcacccccgaaggggac
gacaagcctggcgcagtaggcaaggtggtgcccttcttcgaggctaaggtggtggacttggacaccggtaagacactgggtgtgaaccag
cgcggcgagctgtgcgtccgtggccccatgatcatgagcggctacgttaacaaccccgaggctacaaacgctctcatcgacaaggacggc
tggctgcacagcggcgacatcgcctactgggacgaggacgagcacttcttcatcgtggaccggctgaagagcctgatcaaatacaagggc
taccaggtagccccagccgaactggagagcatcctgctgcaacaccccaacatcttcgacgccggggtcgccggcctgcccgacgacgat
gccggcgagctgcccgccgcagtcgtcgtgctggaacacggtaaaaccatgaccgagaaggagatcgtggactatgtggccagccaggtt
acaaccgccaagaagctgcgcggtggtgttgtgttcgtggacgaggtgcctaaaggactgaccggcaagttggacgcccgcaagatccgc
gagattctcattaaggccaagaagggcggcaagatcgccgtgtaagggtggcatccctgtgacccctccccagtgcctctcctggccctg
gaagttgccactccagtgcccaccagccttgtcctaataaaattaagttgcatcattttgtctgactaggtgtccttctataatattatg
gggtggaggggggtggtatggagcaaggggcaagttgggaagacaacctgtagggcctgcggggtctattgggaaccaagctggagtgca
gtggcacaatcttggctcactgcaatctccgcctcctgggttcaagcgattctcctgcctcagcctcccgagttgttgggattccaggca
tgcatgaccaggctcagctaattttttgtttttttggtagagacggggtttcaccatattggccaggctggtctccaactcctaatctcag
gtgatctacccaccttggcctcccaaattgctgggattacaggcgtgaaccactgctcccttccctgtcctt

**6×cHS4 Insulator** (2143–3576, 1434 bp)
gatccgtcgaccttttccccgtatcccccaggtgtctgcaggctcaaagagcagcgagaagcgttcagaggaaagcgatcccgtgccac
cttccccgtgcccgggctgtccccgcacgctgccggctcggggatgcgggggggagcgccggaccggagcggagccccgggcggctcgctg
ctgcccccctagcgggggagggacgtaattacatccctgggggctttggggggggggctgtccctggcaatcttttccccgtatcccccag
gtgtctgcaggctcaaagagcagcgagaagcgttcagaggaaagcgatcccgtgccaccttccccgtgcccgggctgtccccgcacgctg
ccggctcggggatgcgggggggagcgccggaccggagcggagccccgggcggctcgctgctgccccctagcgggggagggacgtaattaca
tccctgggggctttggggggggggctgtccctcggttttccccgtatcccccaggtgtctgcaggctcaaagagcagcgagaagcgttca
gaggaaagcgatcccgtgccaccttccccgtgcccgggctgtccccgcacgctgccggctcggggatgcgggggggagcgccggaccggag
cggagccccgggcggctcgctgctgccccctagcgggggagggacgtaattacatccctgggggctttggggggggggctgtcccttactt
ttccccgtatcccccaggtgtctgcaggctcaaagagcagcgagaagcgttcagaggaaagcgatcccgtgccaccttccccgtgcccg
ggctgtccccgcacgctgccggctcggggatgcgggggggagcgccggaccggagcggagccccgggcggctcgctgctgccccctagcgg
gggagggacgtaattacatccctgggggctttggggggggggctgtcccttgtttttccccgtatcccccaggtgtctgcaggctcaaag
agcagcgagaagcgttcagaggaaagcgatcccgtgccaccttccccgtgcccgggctgtccccgcacgctgccggctcggggatgcggg
gggagcgccggaccggagcggagccccgggcggctcgctgctgccccctagcgggggagggacgtaattacatccctgggggctttgggg
ggggctgtcccttggttttccccgtatcccccaggtgtctgcaggctcaaagagcagcgagaagcgttcagaggaaagcgatcccgtg
ccaccttccccgtgcccgggctgtccccgcacgctgccggctcggggatgcgggggggagcgccggaccggagcggagccccgggcggctc
gctgctgccccctagcgggggagggacgtaattacatccctgggggctttggggggggggctgtcccttaaagccaccggcaatc

**IRP (CMV minimal)** (3577–3640, 64 bp)
tagagggtatataatggaagctcgacttccagcttggcaatccggtactgttggtaaagccacc

**RRM (rluc2)** (3641–5065, 1425 bp)
gcacctgcaggCatggcttccaaggtgtacgaccccgagcaacgcaaacgcatgatcactgggcctcagtggtgggctcgctgcaagcaa
atgaacgtgctggactccttcatcaactactatgattccgagaagcacgccgagaacgccgtgatttttctgcatggtaacgctacctcc
agctacctgtggaggcacgtcgtgcctcacatcgagcccgtggctagatgcatcatccctgatctgatcggaatgggtaagtccggcaag
agcgggaatggctcatatcgcctcctggatcactacaagtacctcaccgcttggttcgagctgctgaaccttccaaagaaaatcatcttt
gtgggccacgactgggggggctgctctggcctttcactacgcctacgagcaccaagacaggatcaaggccatcgtccatatggagagtgtc
gtggacgtgatcgagtcctgggacgagtggcctgacatcgaggaggatatcgccctgatcaagagcgaagagggcgagaaaatggtgctt
gagaataacttcttcgtcgagaccgtgctcccaagcaagatcatgcggaaactggagcctgaggagttcgctgcctacctggagccattc
aaggagaagggcgaggttagacggcctaccctctcctggcctcgcgagatccctctcgttaagggaggcaagcccgacgtcgtccagatt
gtccgcaactacaacgcctaccttcgggccagcgacgatctgcctaagctgttcatcgagtccgaccctgggttcttttccaacgctatt
gtcgagggagctaagaagttccctaacaccgagttcgtgaaggtgaagggcctccacttcctccaggaggacgctccagatgaaatgggt
aagtacatcaagagcttcgtggagcgcgtgctgaagaacgagcagtaagggtggcatccctgtgacccctccccagtgcctctcctggcc
ctggaagttgccactccagtgcccaccagccttgtcctaataaaattaagttgcatcattttgtctgactaggtgtccttctataatatt
atggggtggaggggggtggtatggagcaaggggcaagttgggaagacaacctgtagggcctgcggggtctattgggaaccaagctggagt
gcagtggcacaatcttggctcactgcaatctccgcctcctgggttcaagcgattctcctgcctcagcctcccgagttgttgggattccag
gcatgcatgaccaggctcagctaattttttgtttttttggtagagacggggtttcaccatattggccaggctggtctccaactcctaatct
caggtgatctacccaccttggcctcccaaattgctgggattacaggcgtgaaccactgctcccttccctgtcctt

### 1 Supplementary Data 1: Complete benchmarking datasets comparing VirDLA predictions with MPRA measurements for tbox__c1 and tbox__h5 in TSV format

**Description:** This dataset contains the complete benchmarking results comparing VirDLA-predicted promoter activities with SHARPR regulatory activity scores. Sequences and scores were extracted from the first 1,000 entries of the SHARPR-MPRA K562 scale-up dataset (minP) (Ernst et al. 2016). The SHARPR scores are deconvolved regulatory activity scores, not raw MPRA RNA/DNA ratios, and were linearized via $2^x$ transformation to align with the VirDLA prediction scale.

**Data files:**

- `Supplementary_Data_1A.tsv` (1,000 rows + header): VirDLA predictions using the **`tbox__c1`** reporter cassette. Columns: `index`, `gt_mpra` (SHARPR regulatory activity score, linearized $2^x$), `pred_vdla` (VirDLA-predicted Fluc/Rluc ratio).
  - Pearson $r = 0.531$, $p = 5.7 \times 10^{-74}$
  - Spearman $\rho = 0.481$, $p = 4.1 \times 10^{-59}$
- `Supplementary_Data_1B.tsv` (1,000 rows + header): VirDLA predictions using the **`tbox__h5`** reporter cassette. Columns: `index`, `gt_mpra` (SHARPR regulatory activity score, linearized $2^x$), `pred_vdla` (VirDLA-predicted Fluc/Rluc ratio).
  - Pearson $r = 0.446$, $p = 5.1 \times 10^{-50}$
  - Spearman $\rho = 0.373$, $p = 2.0 \times 10^{-34}$

### 2 Supplementary Data 2: Pan-tissue activity feature vectors and expression matrices across 273 tissues and cell types in TSV format

**Description:** This dataset contains the pan-tissue VirDLA activity profiles used for the Pan-Tissue Consistency Filter (PTCF) screening. For each of the 10 promoter sequences (WT p16$^{INK4a}$, SRP1M–SRP4M, SRP6M–SRP8M, CMV positive control, Ctrl negative control), VirDLA activity scores (Fluc/Rluc ratio) were predicted across 273 AlphaGenome tissue and cell-type contexts.

**Data file:**

- `Supplementary_Data_2.tsv` (274 lines: header + 273 tissues × 10 promoters):
  - **Header row:** `tissue`, `WT`, `SRP1M`, `SRP2M`, `SRP3M`, `SRP4M`, `SRP6M`, `SRP7M`, `SRP8M`, `CMV`, `Ctrl`
  - Each row represents one tissue context (identified by EFO ontology term or Cell Ontology CL term).
  - Values are VirDLA-normalized Fluc/Rluc activity ratios.
